# VPS28 regulates triglyceride synthesis and is mediated by the ubiquitination pathway in a bovine mammary epithelial cell line and mouse model

**DOI:** 10.1101/2024.07.04.602114

**Authors:** Lily Liu, Jinhai Wang, Xianrui Zheng, Qin Zhang

**Author notes:** Correspondence;, *College of Animal Science and Technology, Shandong Agricultural University, Tai’an 271018, China,* Roslin Institute, University of Edinburgh, Edinburgh EH25 9RG, UK.

## Abstract

VPS28 (vacuolar protein sorting 28) is a subunit of the endosomal sorting complexes required for transport (ESCRTs), and is involved in ubiquitination. Ubiquitination is a crucial system for protein degradation in eukaryotes. Considering the recent findings on the role of ubiquitination in regulating lipid metabolism, we hypothesized that VPS28 might affect the expression of genes involved in milk fat synthesis. To test this hypothesis, we modulated VPS28 expression in the bovine mammary epithelial cell (MAC-T) line and measured the effects on triglyceride (TG) synthesis using lentivirus-mediated techniques. The results indicated that VPS28 knockdown significantly upregulated the fatty acid transporter CD36 (CD36 molecule) and the adipose differentiation-related protein (ADFP), leading to increased TG and fatty acid production, alongside elevated expression of ubiquitin (UB) protein and reduced proteasome activity. In contrast, VPS28 overexpression increased CD36 levels without significantly affecting ADFP and TG levels, showing a trend toward reduced lipid droplets and increased UB expression and proteasome activity. Furthermore, the inhibition of the ubiquitin-proteasome system and endosomal-lysosomal pathway using epoxomicin and chloroquine, respectively, resulted in a further elevation of CD36, ADFP, and TG levels, thereby enhancing cell viability. These in vitro findings were validated in vivo by a mouse model, where VPS28 knockdown enhanced CD36, ADFP, UB expression, TG content, and lipid droplets in mammary glands, without pathological changes in mammary tissue or blood TG alterations. These results confirm the pivotal role of VPS28 in regulating TG synthesis via the ubiquitination pathway, offering novel insights into the molecular mechanisms of milk fat production in a bovine in vitro cell model.

## 1. Introduction

Milk is a rich source of nutrients and consumption of milk is associated with many health benefits. Milk fat is a crucial indicator for evaluating the quality of milk, as it is one of the primary nutrients in milk. It is primarily composed of triglycerides (TG), which serve as a source of energy storage and a means of delivering nutrients, improving taste, and maintaining product structural stability. In the early lactation of the dairy cow, triglyceride synthesis is primarily dependent on de novo fatty acid (FA) synthesis[1]. During peak lactation, absorbed long-chain fatty acids (LFAs) absorbed from blood are predominantly utilized for triglyceride synthesis[1]. Consequently, investigations into the processes, factors, and mechanisms involved in milk fat production in mammary epithelial cells (MECs) are of significant importance for the improvement of milk production and quality.

The synthesis of milk fat in MECs is a physiological reaction associated with multiple steps. These include de novo FA synthesis, FA transport and channeling, TG synthesis, and lipid droplet formation and secretion. A number of molecular players are involved in regulating this process. For instance, the ubiquitination pathway represents a fundamental eukaryotic cellular mechanism, in regulating protein degradation [2]. This ubiquitination pathway can directly regulate the transport of FAs and formation of lipid droplet via CD36 and ADFP [3, 4]. In bovine MECs, CD36 is a key lipid receptor involved in the binding and internalization of a range of lipid species [5, 6]. ADFP is primarily located on the surfaces of lipid droplets, where it is involved in fatty acid synthesis, as well as the transport and exchange of lipids [7].

In our previous study, we found that VPS28 was differentially expressed in mammary tissues from high-and low milk fat dairy cows [8], and there was tissue specific expression of VPS28 in bovine mammary gland, suggesting that VPS28 may play a role in milk fat synthesis [9]. VPS28 is a core component of the class E vacuolar protein -sorting (VPS) proteins, which is essential for integral to the endosomal sorting complexes required for transport (ESCRTs) [10]. The ESCRT system, encompassing the subcomplexes ESCRT-I, -II, and -III, is essential for various cellular undertakings, including nutrient uptake, catabolism, and signaling, by aiding in membrane remodeling and protein transport to lysosomes for degradation[11]. Notably, VPS28 facilitates the recognition of ubiquitinated membrane proteins, and ensuring their proper trafficking and degradation [12, 13]. Available studies have shown that lipid metabolism interacts with vacuoles in yeast and lysosomes in mammalian cells [13, 14]. Previous studies underscore the ESCRT machinery’s role in protein degradation and its disruption leading to lipid accumulation and ubiquitination defects [13–17]. In the mouse, deletion of VPS28 resulted in reduced expression of cell surface proteins [18]. In yeast, VPS28 knockouts fail to deliver ubiquitinated cargo to the vacuole [19]. In humans, disruption of VPS28 resulted in moderate defects in both biosynthetic and endocytic trafficking to the vacuole [20]. Nevertheless, the direct impact of VPS28 on TG and milk fat synthesis in bovine cells remains to be fully explored.

In this sense, the aim of this study was to explore the regulatory mechanisms of VPS28 in milk fat synthesis using bovine mammary epithelial cells (MAC-T) and a mouse model. The anticipated outcomes promise to augment our understanding of the molecular mechanisms of milk fat production.

## 2. Results

### Effects of VPS28 alteration and inhibitors on cell viability in MAC-T cells

The cytotoxicity assay results revealed that neither VPS28 knockdown nor the application of the lysosome inhibitor CQ had a significant impact on the viability of MAC-T cells (*P* > 0.05). In contrast, overexpression of VPS28 markedly reduced cell viability in comparison with the control group, as evidenced by a significant decrease (*P* < 0.01). Additionally, exposure to the proteasome inhibitor EXM resulted in a significant increase in the viability of MAC-T cells (*P* < 0.01) (Figure 1).

**Figure 1.**
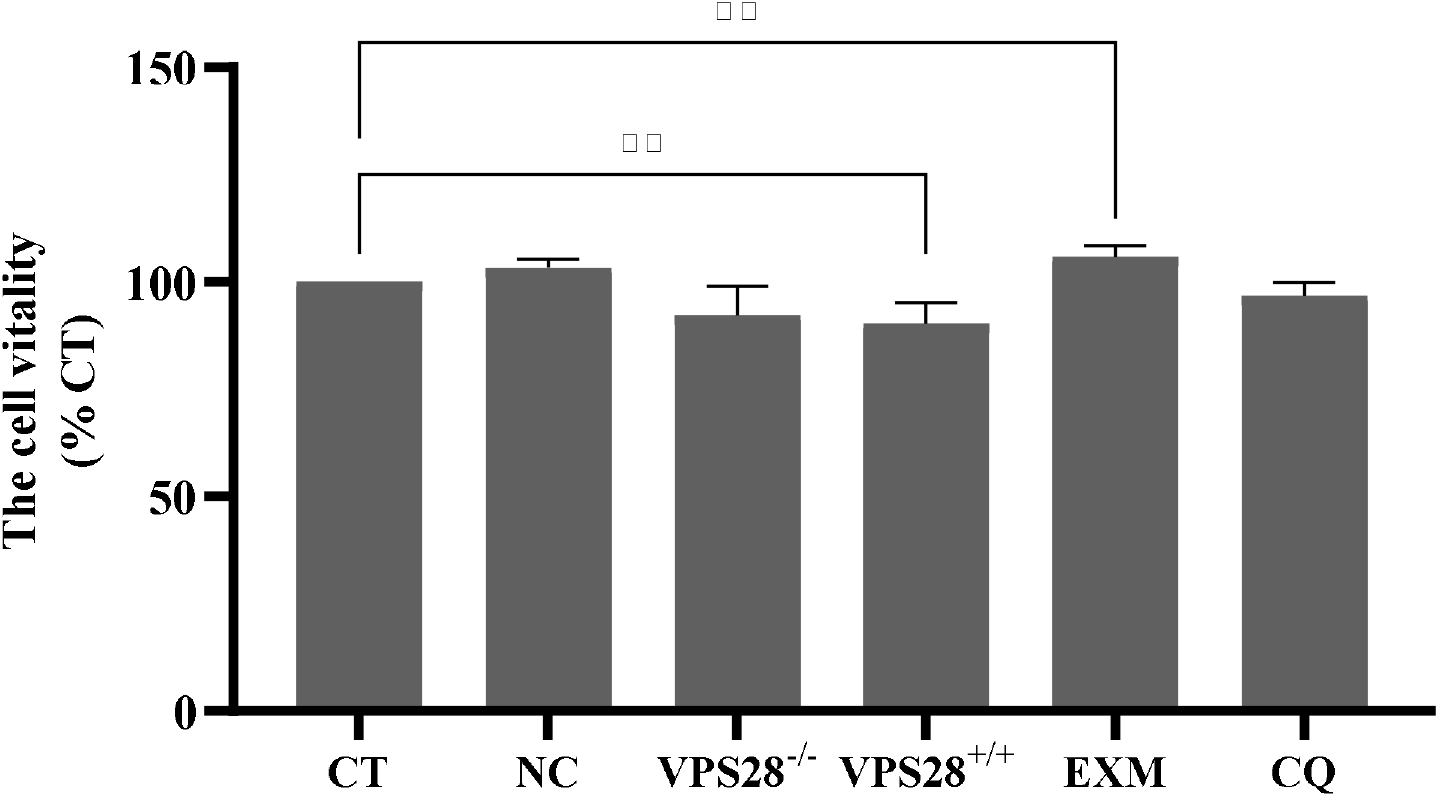
The cell vitality in each group. Treatments were replicated 4 times. The values are means ± SEM. VPS28-/-: VPS28 knockdown. VPS+/+: VPS28 overexpression. ** denote significant differences (P < 0.01, respectively).

### VPS28 knockdown promotes milk fat synthesis in MAC-T cells

In addition, we employed a shRNA sequence targeting VPS28 to suppress its expression in MAC-T cells, which were transfected with the recombinant lentiviral vector LV3. This approach was undertaken to investigate the influence of VPS28 on TG and lipid droplet synthesis. Post-transfection, VPS28 expression was significantly reduced by 26.32% compared to the control group. This reduction was associated with a marked increase in the protein levels of CD36 and ADFP, which increased by 2.53-fold (*P* < 0.05) and 4.51-fold (*P* < 0.01), respectively (Figure 2A). Concurrently, there was a significant enhancement in the TG content of the cells, showing a 1.23-fold increase (*P* = 0.010) (Figure 2B). This increase in TG was accompanied by a significant rise in the number of lipid droplets (Figure 2C).

**Figure 2.**
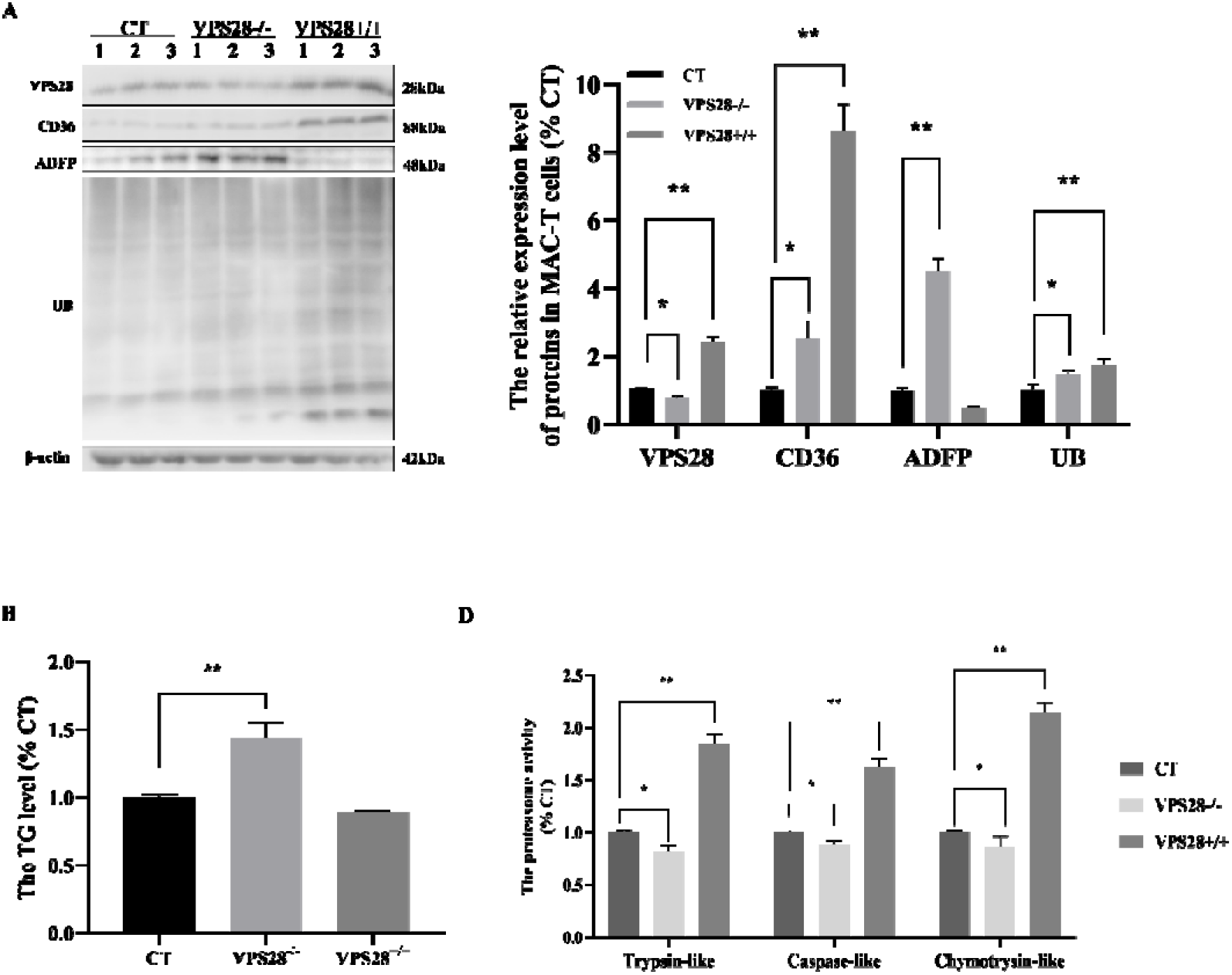
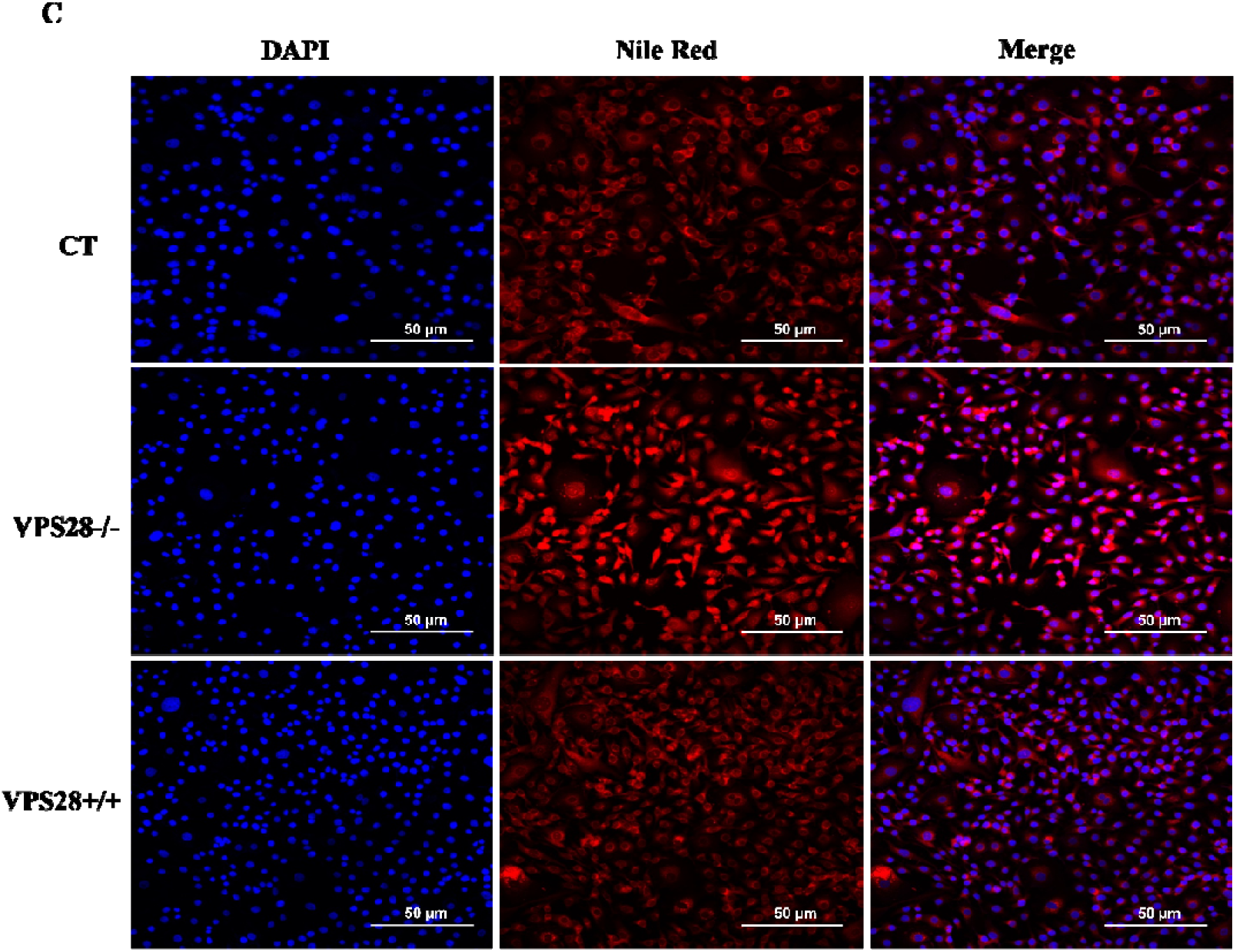
The effects of VPS28 on the MAC-T cells. A: The relative protein expression levels. B: The relative TG content. C: The Nile Red staining. D: The three activities of proteasome. CT: control. VPS28-/-: VPS28 knockdown. VPS+/+: VPS28 overexpression. All treatments were independently replicated three times. Data are presented as means ± SEM. Significance levels: * and ** indicate significant differences (P < 0.05, P < 0.01, respectively).

Moreover, VPS28 knockdown led to a 1.48-fold increase in UB levels (*P* < 0.05). This was accompanied by a decrease of the proteasome’s trypsin-like, caspase-like, and chymotrypsin-like components by 19.04% (*P* < 0.05), 12.17% (P = 0.070), and 14.19% (*P* = 0.482), respectively (Figure 2D). These results underscore the potential of VPS28 knockdown to enhance milk lipid synthesis via the ubiquitination pathway in MAC-T cells.

### VPS28 overexpression promotes ubiquitination pathway in MAC-T cells

To explore the effects of VPS28 overexpression on milk fat synthesis and the dynamics of the ubiquitination pathway, MAC-T cells were transfected with plasmids designed to increase VPS28 expression. This intervention led to a significant elevation in VPS28 levels, with a 2.42-fold increase (*P* < 0.01) (Figure 2A). VPS28 overexpression significantly enhanced CD36 expression by 8.61-fold (*P* < 0.01), while ADFP expression was halved compared to the control group (*P* < 0.05). Despite these changes in gene expression, the concentration of TG remained unaffected (*P* = 0.050), a finding corroborated by Nile Red staining results for lipid accumulation (Figures 2B and C). Furthermore, a significant increase in UB levels was observed, increasing by 1.76 times (*P* < 0.01). Activities of the proteasome components-trypsin-like, caspase-like, and chymotrypsin-like-also saw marked increases of 1.84 (*P* < 0.01), 1.63 (*P* < 0.01), and 2.15 (*P* < 0.01) folds, respectively (Figure 2D). These outcomes indicate that VPS28 overexpression influences milk fat synthesis in MAC-T cells, likely through modulation of the ubiquitination pathway.

### Proteasome inhibitor EXM increased milk lipid synthesis in MAC-T cells

To elucidate the impact of the UPS on milk lipid synthesis within MAC-T cells, we employed the proteasome inhibitor EXM to dampen proteasome activity. Figure 3A shows that proteasome inhibition resulted in reduction in the activities of its trypsin-like, caspase-like, and chymotrypsin-like components, with factors of 0.43, 0.30, and 0.57, respectively (all *P* < 0.01). This inhibition was associated with a 3.06-fold increase in UB expression (*P* < 0.05) (Figure 3B). Moreover, EXM treatment enhanced CD36 and ADFP expression levels by 2.80-fold (*P* < 0.01) and 2.52-fold (*P* < 0.05), respectively.

**Figure 3.**
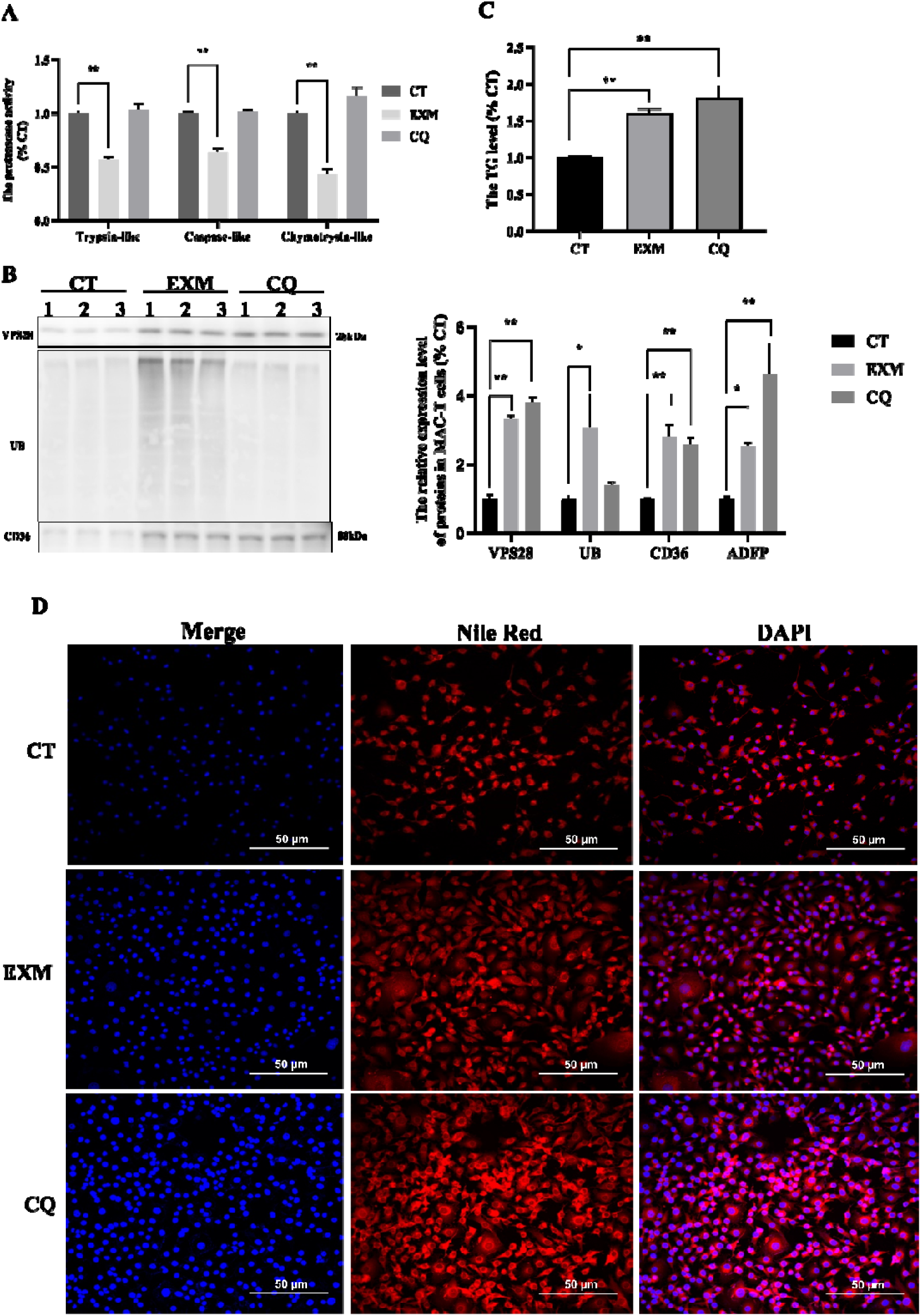
The effect of proteasome and lysosome inhibition in MAC-T. A: The three activities of proteasome. B: The relative protein expression levels. C: The relative TG content. D: The Nile Red staining. CT: control. EXM: inhibited proteasome activity using epoxomicin. CQ: inhibited lysosomal activity using chloroquine. All treatments were independently replicated three times. Data are presented as means ± SEM. Significance levels: * and ** indicate significant differences (P < 0.05, P < 0.01, respectively).

Further analysis involving TG quantification and Nile Red staining (Figures 3C and D) revealed a 1.36-fold augmentation in TG content (*P* < 0.05) coupled with an accumulation of lipid droplets in the MAC-T cells following EXM treatment. These findings underscore a significant link between the UPS and milk fat synthesis in MAC-T cells, highlighting the UPS’s pivotal role in regulating lipid metabolic processes.

### Lysosome inhibitor CQ increased milk lipid synthesis in MAC-T cells

In order to assess the role of ELP in milk lipid synthesis and its contribution to the proteasome-mediated ubiquitination, the CQ inhibitor was employed to impede ELP function in MAC-T cells. The results showed that CQ treatment had no significant impact on the proteasomal or on UB levels.

However, similar to the observations made with EXM, CQ administration resulted in a upregulation of CD36 and ADFP expressions, increasing by 2.58-fold (*P* < 0.01) and 4.63-fold (*P* < 0.01), respectively. Additionally, there was a 1.8-fold elevation in TG content (*P* < 0.01) (Figure 3C), which was accompanied by an increase in lipid droplet accumulation within MAC-T cells (Figure 3D). These findings confirm the significant of role the ELP in regulating TG metabolism in MAC-T cells, highlighting its involvement in modulating milk lipid synthesis through ubiquitination pathways.

### Intraperitoneal injection of VPS28 knockdown and inhibitors does not have a significant influence on the lifespan of mice

In our *in vivo* study, we administered intraperitoneal injections of VPS28 knockdown (using recombinant lentiviral vectors LV3) to mice, along with the proteasome inhibitor EXM and the lysosome inhibitor CQ. This approach aimed to observe the effects of these interventions on body weight (BW) changes. Given the potential risk of VPS28 overexpression inducing apoptosis in MAC-T cells, which could lead to severe inflammation in mice, we opted against creating a VPS28 overexpression mouse model.

Our findings revealed that there were no significant differences in BW alterations among groups (*P* > 0.05) (Figure 4). This outcome showed that the intraperitoneal injection of the lentivirus or the inhibitors did not exert any marked side effects on the mice, indicating the effective establishment of the mouse model for the current study. This result supports the viability of using these interventions *in vivo* without adversely affecting the overall health or BW of the mice, providing a solid foundation for further explorations into the role of VPS28 and ubiquitination pathways in mammalian models.

**Figure 4.**
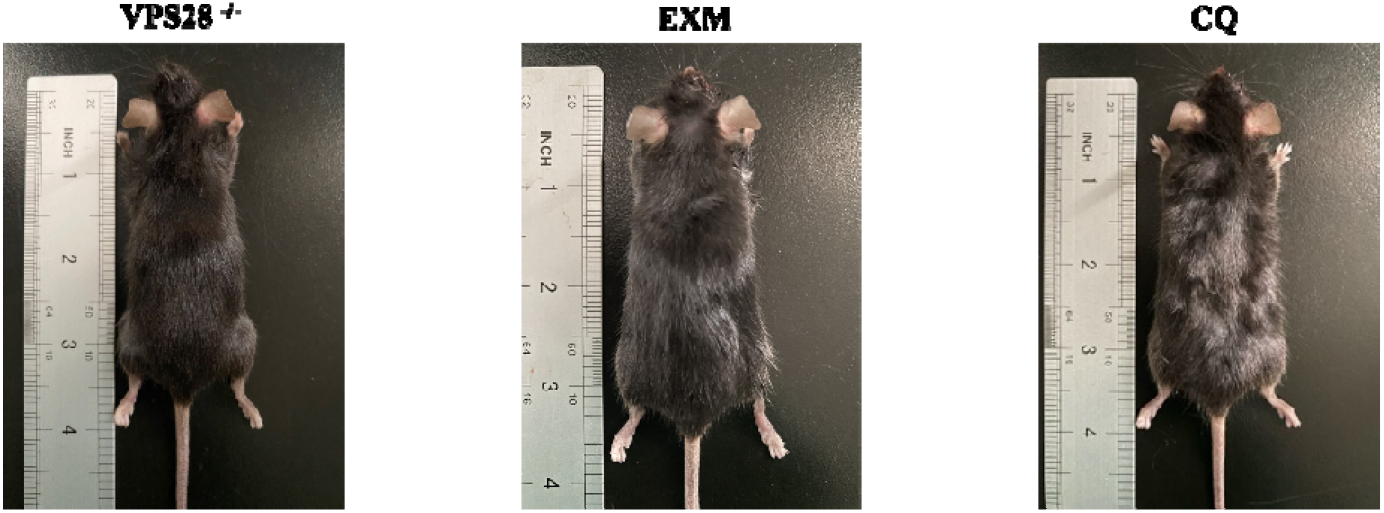
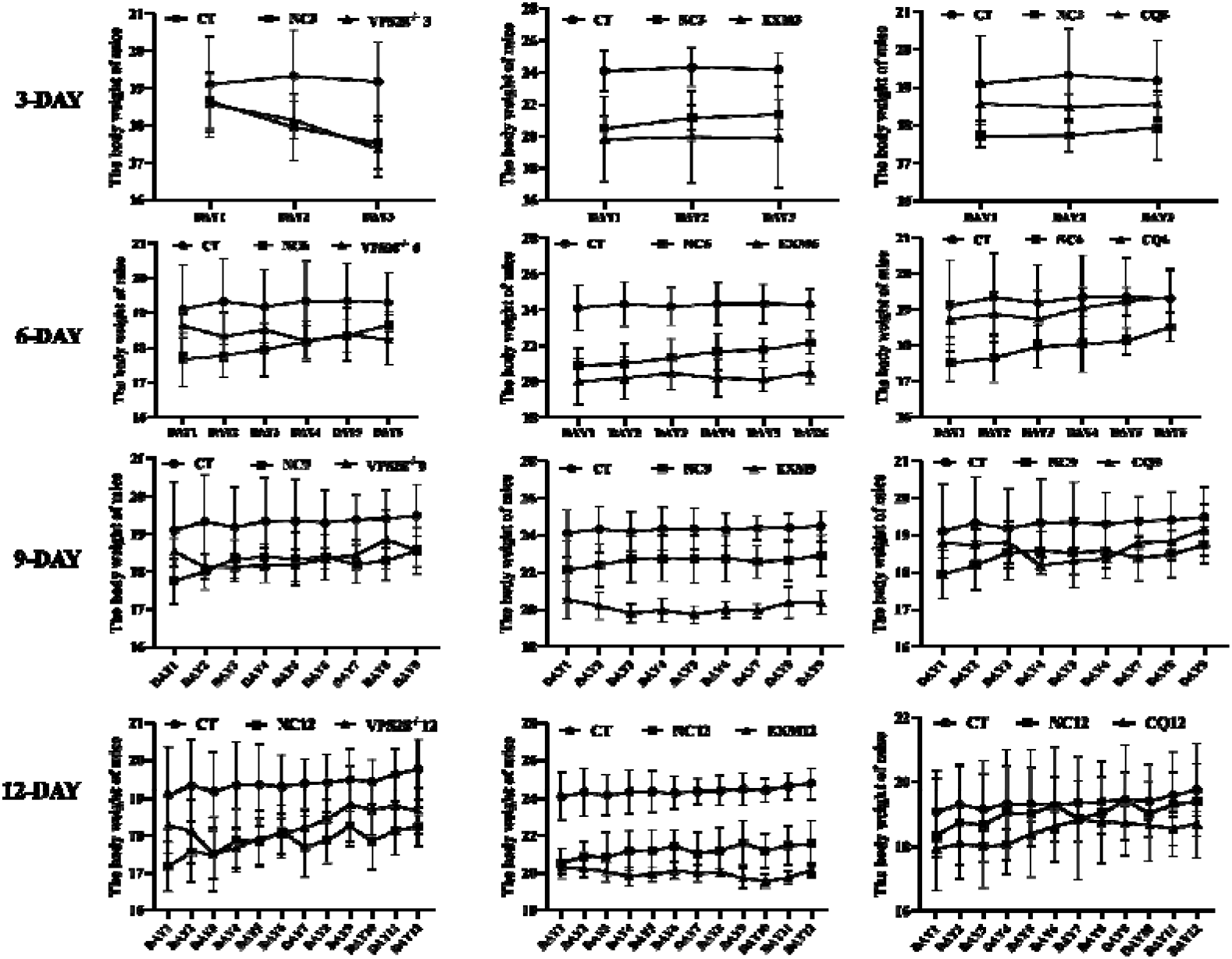
Body weight changes of mice in each group (n=6 per group). Data was collected by weighing each mouse in the treatment groups at regular intervals throughout the study. The error bars represent SD. Statistical significance was determined using a one-way ANOVA followed by post hoc Tukey’s test.

### Intraperitoneal injection of VPS28 knockdown promote lipid synthesis in both mammary gland and blood in mice

After verifying the safety of VPS28 knockdown in mice, we explored its impact on TG synthesis and ubiquitination-related proteins. We first assessed VPS28 protein levels in the mammary glands of mice. The data, depicted in Figures 5A, indicated a marked reduction in VPS28 expression in all treated groups (*P* <0.05), excluding the 6-day cohort, affirming the successful creation of a VPS28 knockdown mouse model via intraperitoneal lentiviral injection. Further analysis of CD36, ADFP, and UB expression levels in mammary gland tissues revealed significant increases compared to control group (*P* <0.01), suggesting that VPS28 knockdown modulates TG synthesis and ubiquitination.

**Figure 5.**
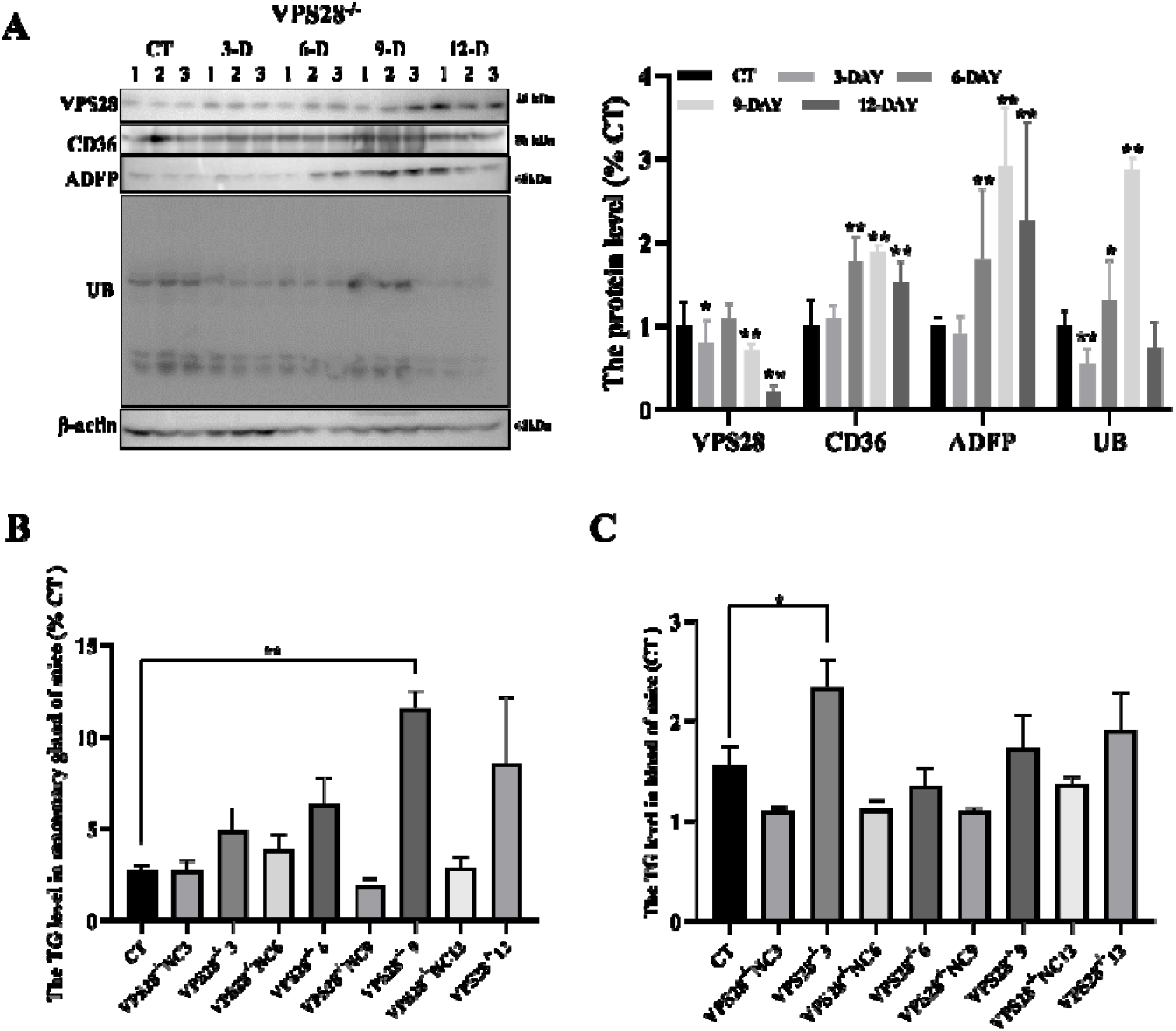
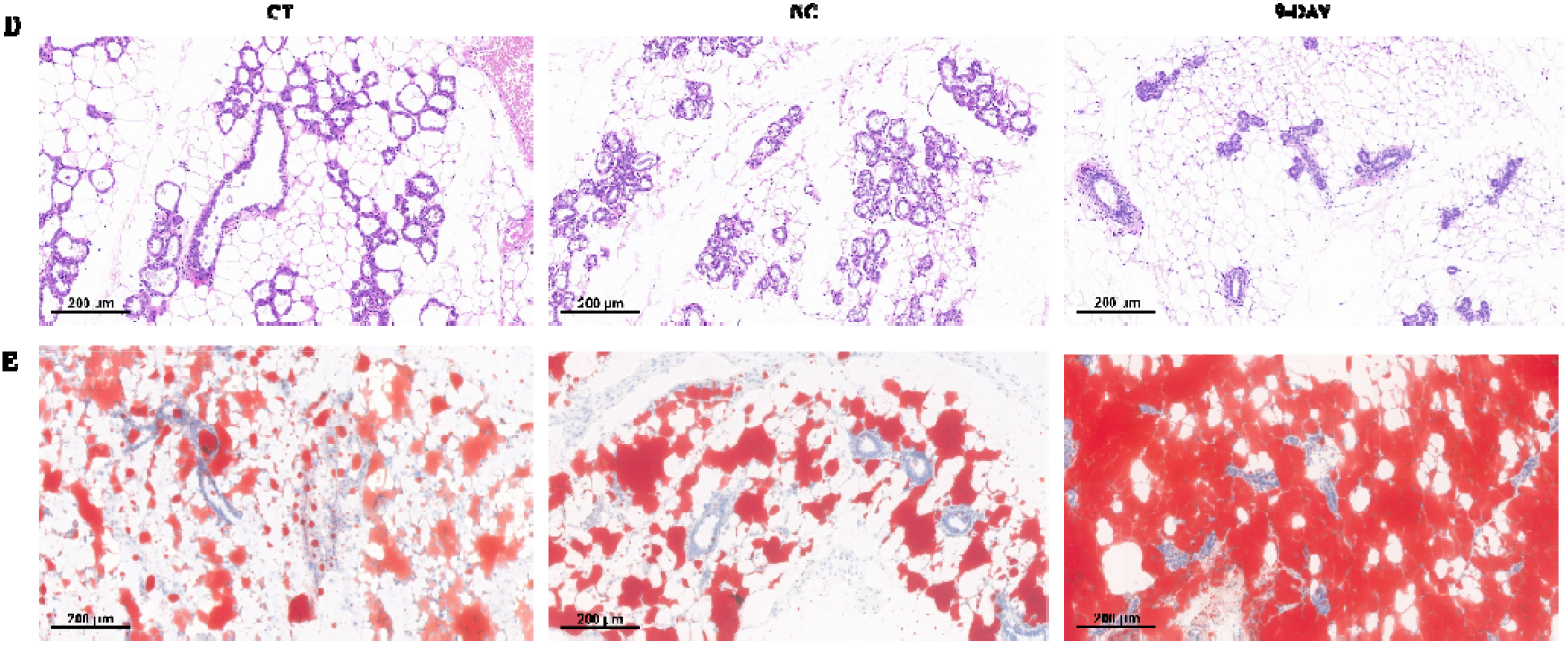
The impact of intraperitoneal injection of VPS28 knockdown on mice. A: Western blot assay was conducted after VPS28 knockdown to assess changes in CD36, ADFP, and UB protein expression in the mammary gland of mice. B-C: The levels of TG in the mammary gland and blood of mice were measured (n=3, error bars represent SEM. Statistical significance was determined using a one-way ANOVA followed by post hoc Tukey’s test). D-E: Representative images illustrating the morphology of the mammary gland using HE staining and Oil Red staining were obtained for each treatment group (n=3 per group).

TG concentrations in mammary glands and blood, along with mammary gland lipid droplet staining, were examined. Notably, VPS28 knockdown elevated mammary gland TG levels, with a peak on day 9 (*P* <0.01), whereas blood TG levels remained largely unaffected, with a minor elevation on day 3 (Figures 5B and C). The 9-day cohort was chosen for detailed lipid droplet analysis in mammary tissues, revealing increased lipid droplet numbers without morphological changes (Figures 5D and E). These results confirm that VPS28 knockdown influences ubiquitination and regulates TG production in the blood and lipogenesis in mammary glands.

### Intraperitoneal Injection of EXM promote lipid synthesis in both mammary gland and blood in mice

In mice treated with EXM intraperitoneally, we observed significant alterations in mammary gland protein levels (Figures 6A). Post-EXM injection, VPS28 expression incrementally rose, while CD36, ADFP, and UB levels initially surged then diminished. Specifically, CD36 decreased significantly on day 12 (*P* <0.05), whereas ADFP saw a substantial increase (*P* <0.01). TG levels in mammary glands and blood were measured, showing a peak in mammary gland TG on day 3 post-injection before decreasing (*P* <0.05), with blood TG levels remaining stable (Figures 6B and C). The 3-day post-EXM treatment model highlighted an increase in mammary gland lipid droplets without tissue morphology alterations (Figures 6D and E), indicating that EXM-mediated inhibition of the ubiquitin-proteasome pathway significantly boosts milk fat production.

**Figure 6.**
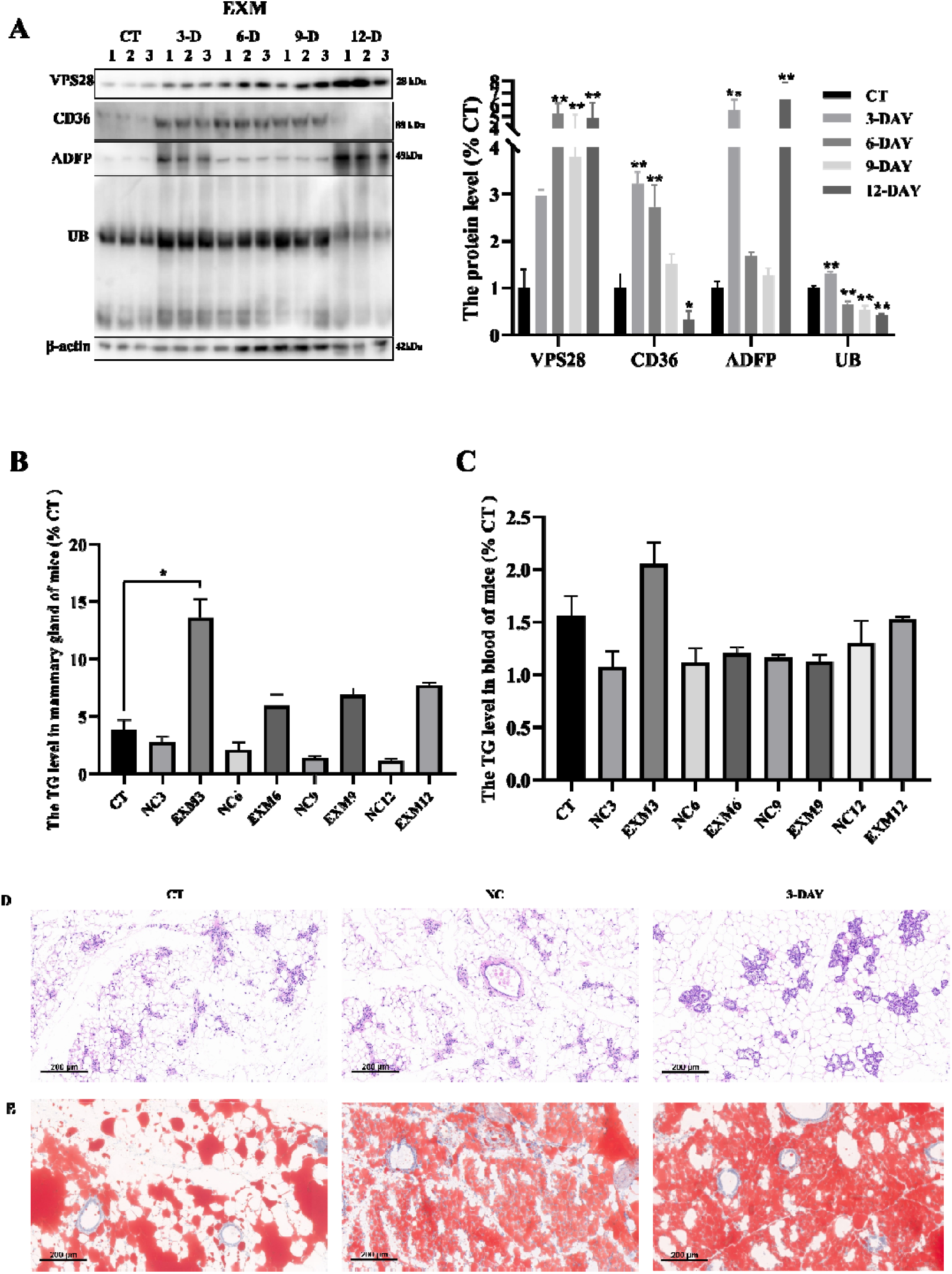
The impact of intraperitoneal injection of EXM on mice was investigated. A: Western blot assay was conducted after EXM injection to assess changes in CD36, ADFP, and UB protein expression in the mammary gland of mice. B-C: The levels of TG in the mammary gland and blood of mice were measured (n=3, error bars represent SEM. Statistical significance was determined using a one-way ANOVA followed by post hoc Tukey’s test). D-E: Representative images illustrating the morphology of the mammary gland using HE staining and Oil Red staining were obtained for each treatment group (n=3 per group).

### Intraperitoneal Injection of CQ promote lipid synthesis in both mammary gland and blood in mice

Similarly, in the CQ-treated mice, significant protein level changes were noted in mammary gland tissues post-injection (Figures 7A). Following CQ treatment, VPS28 expression significantly increased (*P* < 0.01), and CD36 (*P* < 0.05), ADFP (*P* < 0.01), and UB (*P* < 0.01) levels notably rose on days 9 and 12. TG measurement results showed a peak in mammary gland TG levels on day 9 post-injection (*P* < 0.01), with stable blood TG levels (Figures 7B and C). The 9-day CQ treatment model exhibited a significant rise in mammary gland lipid droplets, maintaining normal tissue morphology (Figures 7D and E). This suggests that inhibiting the ubiquitin-lysosome pathway with CQ effectively enhances milk fat production in mouse mammary glands.

**Figure7.**
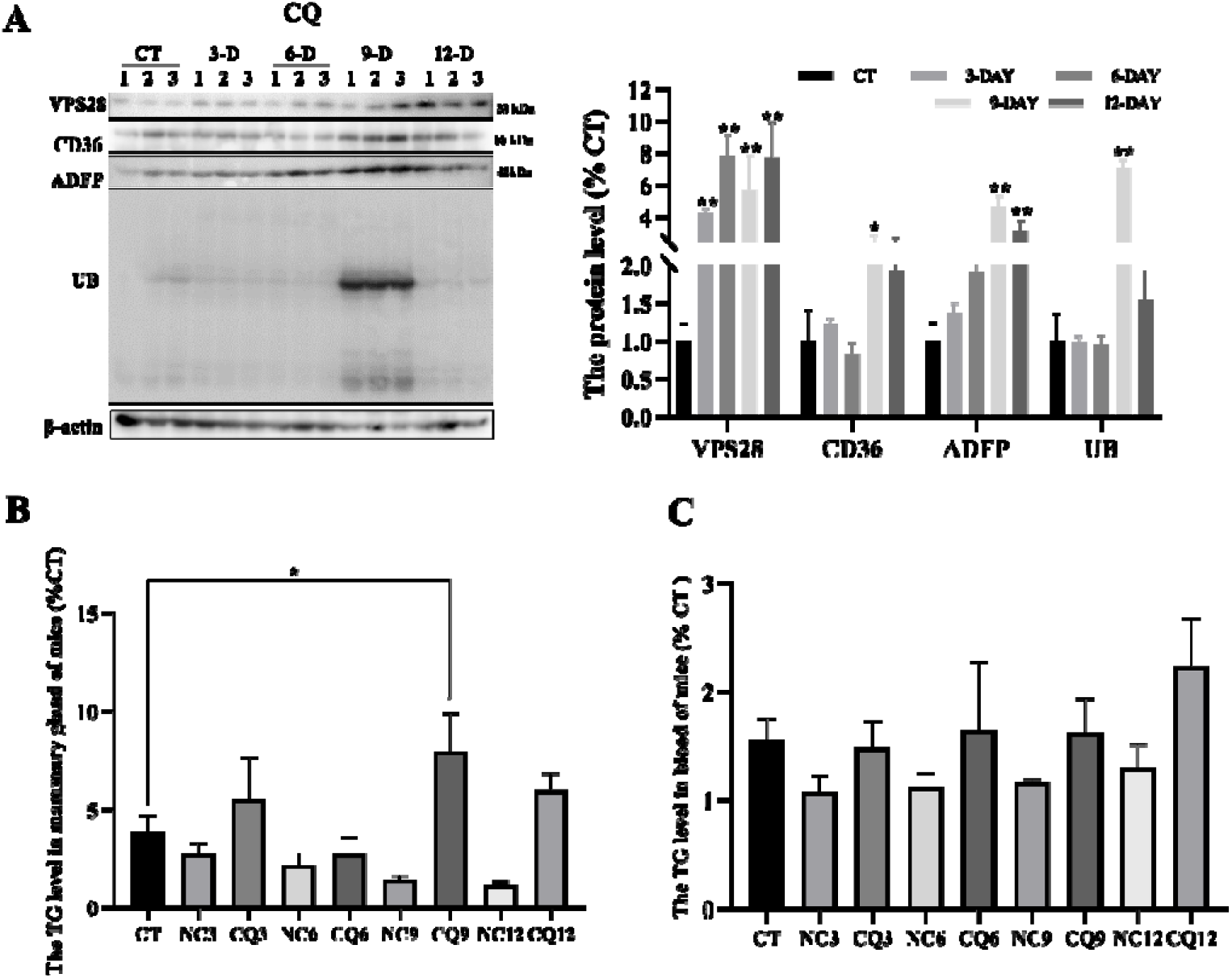
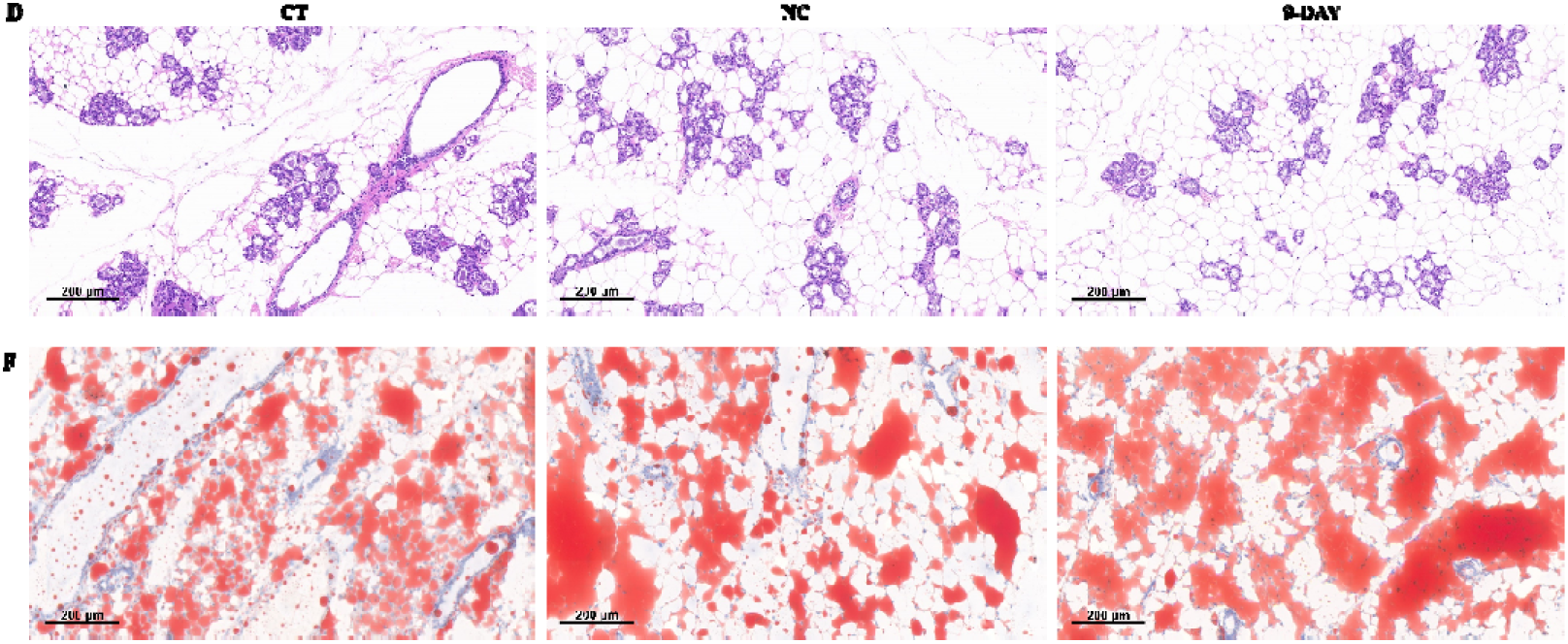
The impact of intraperitoneal injection of CQ on mice was investigated. A: Western blot assay was conducted after CQ injection to assess changes in CD36, ADFP, and UB protein expression in the mammary gland of mice. B-C: The levels of TG in the mammary gland and blood of mice were measured (n=3, error bars represent SEM. Statistical significance was determined using a one-way ANOVA followed by post hoc Tukey’s test). D-E: Representative images illustrating the morphology of the mammary gland using HE staining and Oil Red staining were obtained for each treatment group (n=3 per group).

## 3. Discussion

In our previous study, we found there was a C-58T mutation in *VPS28* gene, that could lead a reduction expression in VPS28 gene and was significantly associated with high milk fat percentage in Holstein dairy cows [9]. Therefore, we hypothesis that the lower expression of VPS28 may facilitate milk fat synthesis in bovine. To validate it, we conducted the identification in MACT cell lines and mouse model.

VPS28, an integral component of the ESCRT complex, plays a pivotal role in cellular processes such as endocytosis, protein sorting, and the degradation of membrane proteins [19, 21] [22]. The ESCRTs contain multiple members involved in endosomal-lysosomal pathway (ELP), regulating ubiquitin-mediated degradation of membrane proteins, crucial for the balance of receptor-mediated signaling pathways [23–28]. VPS28 could also affect the function of the ubiquitin-proteasome system (UPS) [29–31], which is the major regulator of all ubiquitinated proteins in cells [32–34]. Recent studies suggest that ubiquitination plays a much broader role in regulating protein function for regulating levels of transporters or some membrane proteins [35–37]. The process of ubiquitination was observed to be paralleled by alterations in the uptake and lipid incorporation of the FA in human and rat cells [38, 39]. Furthermore, VPS28 was identified as playing a vital role in the regulation of growth and the cell cycle in muscle cells of sheep [40], and has been demonstrated to regulate hair follicle differentiation in goats [41].

In order to further elucidate the role of VPS28 in milk fat synthesis, we conducted knockdown and overexpression experiments in VPS28 in MAC-T cells and assessed the effects on milk fat synthesis by measuring the expression levels of the lipid synthesis-related genes CD36 and ADFP, TG content and lipid droplet content. The results showed that lower expression of VPS28 promoted the expression of the CD36 and ADFP, and TG content in MAC-T cells. VPS28 can regulate the accumulation of ubiquitinated protein levels through the ELP and UPS. Therefore, we hypothesis that VPS28 may regulate milk fat synthesis in MAC-T cells by altering the ELP and UPS pathways. The experimental results corroborated our hypothesis that VPS28 knockdown resulted in an accumulation of UB levels and an inhibition of proteasome. Conversely, corresponding overexpression of VPS28 was found to promote the proteasome activity. It has been demonstrated that ubiquitinated CD36 facilitates the transport FAs into cells, thereby promoting TG synthesis and the formation of lipid droplets [42–45]. ADFP is a protein involved in lipid management and is essential for lipid metabolism and storage within cells [7]. It is also regulated by the ubiquitination pathway [46, 47]. This outcome provides evidence that the alteration of VPS28 directly affects the level of ubiquitinated CD36 and ADFP, and regulates TG and lipid droplet synthesis in MAC-T cells through the ubiquitination process.

In consideration of the relationship among VPS28, ELP and USP, we introduced the lysosomal inhibitor CQ and proteasomal inhibitor EXM to MAC-T cells. CQ, a medication commonly used to treat malaria and autoimmune conditions, impedes the lysosomal fusion process and modifies the acidic environment within lysosomes [48]. This disruption affects the distribution and breakdown of substances within cells, as evidenced by previous study [49]. Moreover, studies indicate that CQ may influence autophagy, a cellular degradation mechanism, potentially affecting the removal and processing of cellular debris [50]. EXM targets key proteins within proteasomes, particularly binding to the active sites of the 20S proteasome, thus obstructing the proteasomal degradation route and hindering the breakdown of ubiquitinated proteins within cells [51, 52]. It has been demonstrated that the inhibition of ubiquitination signaling pathways can lead to the accumulation of ubiquitinated CD36 and ADFP [42, 53]. This accumulation could be the results of post-translational regulation by the ubiquitination pathway, particularly in the context of proteasome inhibition. In MAC-T cells, the results demonstrated that both CQ and EXM did not affect the cell viability and could increase the expression of VPS28, CD36, and ADFP. This was accompanied by a significant rise in TG content and lipid droplet accumulation. It is speculated that these effects may be caused by the ubiquitination pathways mediated by ELP and USP, which underscores the linkage among VPS28, the ELP and UPS.

Furthermore, we inhibited VPS28 expression by intraperitoneal injection of lentivirus and demonstrated *in vivo* in mice that knockdown of VPS28 promoted milk fat synthesis and the ubiquitination level. These outcomes provide compelling evidence to support our hypothesis that VPS28 directly influences TG and lipid droplet synthesis in MAC-T cells through the ubiquitination process. Moreover, we employed CQ and EXM to inhibit lysosomal and proteasomal activity. The effects of CQ and EXM treatment revealed no adverse effects on the liver and mammary gland tissues of mice, indicating the treatment’s safety at the utilized concentrations. Concomitantly, there was a marked increase in the levels of VPS28, CD36, and ADFP. Notably, following CQ and EXM treatment, there was a pronounced increase in TG levels and lipid droplet accumulation, exceeding the effects observed when VPS28 expression was modified or CQ or EXM administered. This finding not only verifies VPS28’s regulatory capacity over TG and lipid droplet synthesis via the UPS but also highlights the UPS’s central role in TG and lipid droplet production.

## 4. Conclusions

The present study demonstrated that, in MAC-T cells, VPS28 knockdown promotes milk fat synthesis by regulating the ubiquitination pathways and targeting ubiquitinated CD36 and ADFP. This study elucidated the function of VPS28 in milk fat synthesis and identifies the potential utility of EXM in promoting milk fat synthesis. The results of this study have contributed to a greater understanding of the regulatory network of milk fat synthesis using a bovine *in vitro* system.

## 5. Materials and Methods

### Reagents

CQ (CAS No. C6628-25G) was purchased from Sigma (Sigma-Aldrich, USA), EXM (CAS No. 134381-21-8) and Nile Red (CAS No. 7385-67-3) were purchased from Med Chem Express (MCE, USA). Antibodies were purchased from Abcam (Cambridge, UK). All other reagents were purchased from Life Technologies (Carlsbad, CA, USA) unless noted otherwise. EXM and CQ were dissolved in a solution comprising 10% DMSO and 90% saline with 20% SBE-β-CD to use.

### Cell culture and treatments

The MAC-T cells were first cultured in serum-containing medium DMEM, supplemented with 10 kU/mL penicillin, 10mg/mL streptomycin, and 10% FBS (fetal bovine serum). All cells were cultured in plastic cell culture plates at 37L in a humidified atmosphere containing 75% CO_2_. VPS28 was knocked down using recombinant lentiviral vector LV3 (H1/GFP&Puro) containing RNA interference fragments (shRNA: CCGGGGACGTGGTCTCGCTCTTTATCTCGAGATAAAGAGCGAGACCACGTCCTTTTTG), and overexpressed using recombinant lentiviral vectors pCDH-CMV-MCS-EF1-GFP-Puro-VPS28. For lentivirus-mediated overexpression, MAC-T cells cultured in six-well plates were incubated with lentiviral helper plasmids PSPAX2 and pMD2.G, while the control was treated with lentivirus encapsulated in an empty vector. All lentiviruses were ensured to have a lentiviral titer of 2×10^8 plaque-forming units per mL. After 72 hours, cells were harvested for protein collection.

Proteasome-inhibited MAC-T cells were cultured with 1 μmol/L EXM. Lysosome-inhibited MAC-T cells were cultured with 50 μmol/L CQ. Control cells were treated with solution of EXM and CQ. Following a 24-hour incubation, all cells were collected for protein extraction, as well as analysis of TG.

### Mice

The in vivo experiments were performed with 150 female mice, 7-8 weeks old, accommodated at the Kunming Institute of Zoology, Chinese Academy of Sciences, located in Kunming, China. The animals were kept under controlled environmental conditions at a temperature of 25°C with a humidity level set at 55%. They were provided with 12 h of ambient lighting in ventilated and pathogen-free cages. The animals had free access to drinking water and food and their health status was checked daily. The ethical approval for the use and care of animals in this study was granted by the Academic Committee of Southwest Forestry University (Approval ID: SWFU-20220307-1). We confirm that all experimental procedures and protocols conducted in this study were performed in accordance with the relevant guidelines and regulations, including the ARRIVE guidelines for reporting animal research. The study protocol was approved by the Academic Committee of Southwest Forestry University.

### Treatments

The mice were randomly assigned to a control group (n=6), VPS28 knockdown group (n=48), EXM model group (n=48) and CQ model group (n=48). The three model groups were established by continuous intraperitoneal injections of (100 μL per day of VPS28-knockdown lentivirus with a lentiviral titer of 2×10^8 plaque-forming units per mL; 5 μM per day of EXM with a concentration of 1 mg/mL; 3.1 mM per day of CQ with a concentration of 1.6 mg/mL) for 12 days. Afterward, every model group were randomly divided into 8 smaller groups (n=6 each): 3-day group (n=6 each), 6-day (n=6 each), 9-day (n=6 each) and 12-day (n=6 each) and corresponding negative control (NC) group (n=6 each). The NC for EXM and CQ groups were treated with a solution containing 10% DMSO and 90% saline supplemented with 20% SBE-β-CD, and the NC for the VPS28 knockdown group was treated with an empty viral vector. Throughout the study, all mice were provided with a normal chow diet (SWS9102-1010086, Jiangsu Synergetic Pharmaceutical Bioengineering Co., LTD, China) and had access to drinking water.

Throughout the experiment, the body weight (BW) of the mice was meticulously recorded daily for the entire study period to monitor any fluctuations resulting from administration, following the methodology established in a previous study, as a means to assess mice growth performance parameters.

Following injection, blood samples were collected from the eyes of six mice per group for sample collection. Subsequently, the mice were euthanized by cervical dislocation, and samples of the mammary gland were obtained post-sacrifice. For histomorphology analysis, mammary gland from three mice per group were fixed in a 4% polyformaldehyde solution for 24 hours.

The remaining tissues from the other three mice per group were washed with ice-cold PBS for the analysis of TG content. Furthermore, all other tissues were collected and stored at −80°C for subsequent western blot analysis. The methods are consistent with the method used for the cell samples.

### Western Blot analysis

After being treated differently according to experimental assignments, MAC-T cells or mammary gland from mice were lysed in cold lysis buffer at 4 °C for 30 min, broken by ultrasonic waves, and centrifuged at 12, 000 × g for 20 minutes. The supernatants were collected, and the protein was quantified using a BCA protein assay kit (Beyotime Biotechnology, China). VPS28 (No.ab154793, 1:500), CD36 (No.ab252922, 1:1,000), ADFP (No.ab181452, 1:1,000) and UB (No.sc-53509, Santa Cruz, USA, 1:500) levels were detected by western blotting, with anti-β-actin (No.66009, Proteintech Group, USA, 1:1,000) used as an internal control. The bands were detected using a chemiluminescent ECL system (Promega, USA) and quantified with Image J (version 1.8.0).

### TG content analysis

To analyze TG content, cellular and tissue samples underwent extraction and quantification using a TG assay kit (CAS: A110-1-1, Nanjing Jiancheng Bioengineering Institute, China) according to the method described in a previous study[54]. Briefly, after adding PBS, the cells were subjected to ultrasonic disruption for 300 seconds, followed by the addition of a working solution for a 30-minute reaction. Subsequently, the absorbance was measured using an enzyme-linked immunosorbent assay (ELISA) reader at a wavelength of 510nm. Concurrently, the total protein concentration in the cells or tissues was determined using BCA, and the TG content in the cells and tissues were calculated using the following formula.

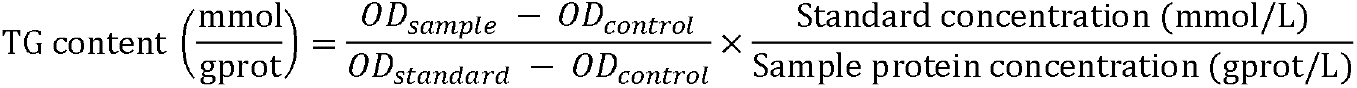

### Nile Red staining

The Nile Red staining was also performed on MAC-T cells according to the manufacturer’s protocol. The cells were seeded into 12-well plates and cultured for 24 hours. After that, the cells were fixed with 4% paraformaldehyde for 15 minutes at room tem-perature and washed three times with 1% BSA in PBS. Subsequently, the cells were stained with 10 µg/mL Nile Red at 4°C for 15 minutes and washed three times with 1% BSA in PBS. The stained cells were then incubated with DAPI (No.P0131, Beyotime Bio-technology, China) until air-dried naturally. Finally, the cells were covered with a cover slip and the lipids were visualized using a confocal microscope (Olympus IX81-FV1000) to capture images.

For Nile Red staining of tissue, fresh mammary gland tissue was fixed in a 4% paraformaldehyde solution for 24 hours. Subsequently, the tissue was embedded in OCT (Optimal Cutting Temperature) and 8-μm-thick sections were obtained for staining. The protocol used for the tissue samples is consistent with the method used for the cell samples.

### Oil Red O staining

Frozen sections, 8 μm thick, were utilized for Oil Red O staining. The staining process involved preparing a 0.5-5% (w/v) Oil Red O working solution by dissolving the powder in isopropanol. Tissue sections were then incubated in this solution for 10-20 minutes at room temperature, followed by a brief rinse in distilled water to remove excess stain. Microscopic observation of mammary gland alveoli and fat deposition was conducted in accordance with the protocol.

### Hematoxylin-eosin (HE) staining

The fresh mammary gland tissue was fixed in a 4% paraformaldehyde solution for 24 hours. Subsequently, the tissue was embedded in paraffin, and 8-μm-thick sections were obtained and subjected to staining using the following steps: 1) Dewaxing: The sections were treated with xylene for 5 minutes and repeated thrice, followed by rinsing with anhydrous ethanol for 5 minutes and repeating the rinse thrice. 2) Ethanol Gradient Elution: The sections were sequentially washed with 95%, 80%, 75% ethanol, and distilled water for 2 minutes each. 3) Hematoxylin Staining: The sections were stained with hematoxylin drops for 5 minutes, followed by rinsing with tap water for 2 minutes. 4) Differentiation: The sections were differentiated with 1% hydrochloric acid ethanol for 30 seconds, followed by soaking in antiblue treatment for 15 minutes. 5) Eosin Staining: The sections were stained with Kay red dye drops for 2 minutes, followed by rinsing with tap water for 3-5 seconds. 6) Dehydration: Dehydration of the sections was carried out using 70% ethanol, 80% ethanol, 95% ethanol, 100% ethanol (twice), and xylene (thrice), each for 1 minute. 7) Sealing: Finally, neutral resin was applied for sealing. Microscopic examination and image acquisition were performed using an orthotopic light microscope (Nikon Eclipse E100, Nikon, Japan) equipped with an imaging system (Nikon DS-U3, Nikon, Japan).

Photomicrographs of three sections per mouse were captured at a magnification of 200×.

### Proteasome activity analysis

The proteasome activities were analyzed by measuring cells using the Proteasome-Glo™ Chymotrypsin-Like, Caspase-Like, and Trypsin-Like Cell-Based Assays (No. G1180, Promega, USA), following the method proposed in a previous study [54]. Briefly, the cells were lysed in a buffer containing a proteasome inhibitor and the lysates were added to the substrate mixtures supplied in the kit. The mixtures contained luminogenic peptide substrates specific for the chymotrypsin-like, caspase-like, or trypsin-like activity of the proteasome. After incubation at room temperature for 15 minutes, the luminescence was measured using a microplate reader.

### Cell viability analysis

The CCK-8 method was utilized for the cell viability assay. MAC-T cells were seeded in a 96-well cell culture plate and allowed to reach 70% confluence. Subsequently, the cells were treated with VPS28 knockdown, VPS28 overexpression, EXM solution, and CQ solution as described above. Following the corresponding transfection and incubation period, the culture medium was replaced with 10% CCK-8 basic medium, and the cells were further incubated for 3 hours. The absorbance of each cell group was measured at a wavelength of 450 nm using an automated microplate reader. Cell viability was determined based on the absorbance values obtained from the assay.

## Statistical analysis

Each treatment was performed in triplicate, and the results were expressed as means ± standard error of the means (SEM). The densitometry values of western blots were measured using Image J software (version 1.8.0), and protein abundance was normalized to β-actin. The data were analyzed using analysis of variance and multiple testing with SPSS Statistics software (version 22.0), and graphs were performed using GraphPad Prism 8.0.1 software (GraphPad, San Diego, CA, USA). Differences were considered significant at *P* < 0.05 and highly significant at *P* < 0.01.

## Acknowledgments

The authors express their gratitude to the Chinese Academy of Sciences, Kunming Institute of Zoology, and the Roslin Institute, University of Edinburgh, for their technical assistance and platform support.

## Author contributions

Conceptualization, L.L. and Q.Z.; methodology, L.L.; software, L.L.; formal analysis, L.L.; investigation, L.L.; resources, L.L; data curation, L.L.; writing—original draft preparation, L.L.; writing—review and editing, L.L., J.W., E.C. and Q.Z.; visualization, L.L.; supervision, L.L., J.W., E.C. and Q.Z.; project administration, L.L., X. Z.; funding acquisition, L.L. All authors have read and agreed to the published version of the manuscript.

## Abbreviations

VPS28: Vacuolar Protein Sorting 28
MAC-T: Bovine Mammary Epithelial Cell Line
CD36: Fatty Acid Transporter CD36
ADFP: Adipose Differentiation-Related Protein
TG: Triglyceride
UB: Ubiquitin
UPS: Ubiquitin-Proteasome System
ELP: Endosomal-Lysosomal Pathway
ESCRT: Endosomal Sorting Complexes Required for Transport
EXM: Epoxomicin
CQ: Chloroquine
HE: Hematoxylin-eosin
BW: body weight
FBS: Fetal Bovine Serum
NC: negative control
WB: Western Blot
OCT: Optimal Cutting Temperature.

## Data Availability Statement

All data measured or analyzed during this work are available from the corresponding author upon reasonable request.

